# Inferring the molecular mechanisms of noncoding Alzheimer’s disease-associated genetic variants

**DOI:** 10.1101/401471

**Authors:** Alexandre Amlie-Wolf, Mitchell Tang, Jessica Way, Beth Dombroski, Ming Jiang, Nicholas Vrettos, Yi-Fan Chou, Yi Zhao, Amanda Kuzma, Elisabeth E. Mlynarski, Yuk Yee Leung, Christopher D. Brown, Li-San Wang, Gerard D. Schellenberg

**Author notes:** To whom correspondence should be addressed. Mailing address: Richards Building, D 101. 3700 Hamilton Walk. Philadelphia, PA 19104. These authors contributed to the manuscript equally (Alexandre Amlie-Wolf, Mitchell Tang, Li-San Wang, Gerard D. Schellenberg). **Declarations of interest:** none.

## Abstract

**INTRODUCTION:** We set out to characterize the causal variants, regulatory mechanisms, tissue contexts, and target genes underlying noncoding late-onset Alzheimer’s Disease (LOAD)-associated genetic signals.

**METHODS:** We applied our INFERNO method to the IGAP genome-wide association study (GWAS) data, annotating all potentially causal variants with tissue-specific regulatory activity. Bayesian co-localization analysis of GWAS summary statistics and eQTL data was performed to identify tissue-specific target genes.

**RESULTS:** INFERNO identified enhancer dysregulation in all 19 tag regions analyzed, significant enrichments of enhancer overlaps in the immune-related blood category, and co-localized eQTL signals overlapping enhancers from the matching tissue class in ten regions (*ABCA7, BIN1, CASS4, CD2AP, CD33, CELF1, CLU, EPHA1, FERMT2, ZCWPW1*). We validated the allele-specific effects of several variants on enhancer function using luciferase expression assays.

**DISCUSSION:** Integrating functional genomics with GWAS signals yielded insights into the regulatory mechanisms, tissue contexts, and genes affected by noncoding genetic variation associated with LOAD risk.

## 1. Background

Alzheimer’s disease (AD) is the most common cause of dementia in the United States [1], but no effective therapies for treatment or prevention exist. Late-onset Alzheimer’s disease (LOAD), defined by age-at-onset after 60 years, is the most common form of AD. Heritability estimates for LOAD stand at 60-80%, implicating genetics as an important factor in disease development [2]. While the *APOE* locus shows the strongest association [3], LOAD is complex and polygenic [4], and genome-wide association studies (GWAS) have successfully associated over 20 other genetic variants with LOAD [5,6]. Recent studies have implicated a number of different biological processes in LOAD susceptibility such as microglial-mediated innate immunity [7–9].

The majority of top GWAS variants reside in noncoding regions of the genome outside of protein-coding sequences [10]. Any variant in linkage disequilibrium (LD) with a top GWAS variant could be responsible for the association signal, and GWAS data alone lacks the granularity to identify these causal variants. In addition, noncoding variants presumably affect gene regulatory elements, and the affected target genes are often not the closest ones [11]. Thus, functional annotation is needed to reveal the causal variants, regulatory mechanisms, tissue context, and target genes underlying GWAS signals.

Enhancers, which modulate the expression of a target gene independently of orientation and distance, are one of the most common regulatory elements in the noncoding genome [12–14]. Several consortia have generated large-scale functional genomics datasets to characterize regulatory activity in the noncoding genome across different tissue contexts [15–19]. Previous studies used these data to identify noncoding genetic variants with regulatory potential for diabetes [20,21] and schizophrenia [22], but such studies often assume that the relevant tissue context is known *a priori*.

We hypothesize that noncoding LOAD GWAS signals modulate disease risk by perturbing genomic elements that regulate genes involved in pathogenesis. To explore this hypothesis, we applied our bioinformatics pipeline, INFERNO (INFERring the molecular mechanisms of NOncoding genetic variants) [23] to LOAD GWAS data from the International Genomics of Alzheimer’s Project (IGAP) [6]. INFERNO characterizes noncoding GWAS signals by integrating information across diverse functional genomics data sources to identify causal noncoding variants and the regulatory mechanisms, tissue contexts, and target genes they affect (Figure 1a). INFERNO identified several putatively causal genetic variants in ten GWAS regions and uncovered strong functional evidence of their effects on immune- and brain-related regulatory mechanisms. Using luciferase reporter assays, we validated the enhancer activity and allelic differences of causal variants in three regions prioritized by relevant tissue context, strength of annotation support, and prior literature.

**Figure 1:**
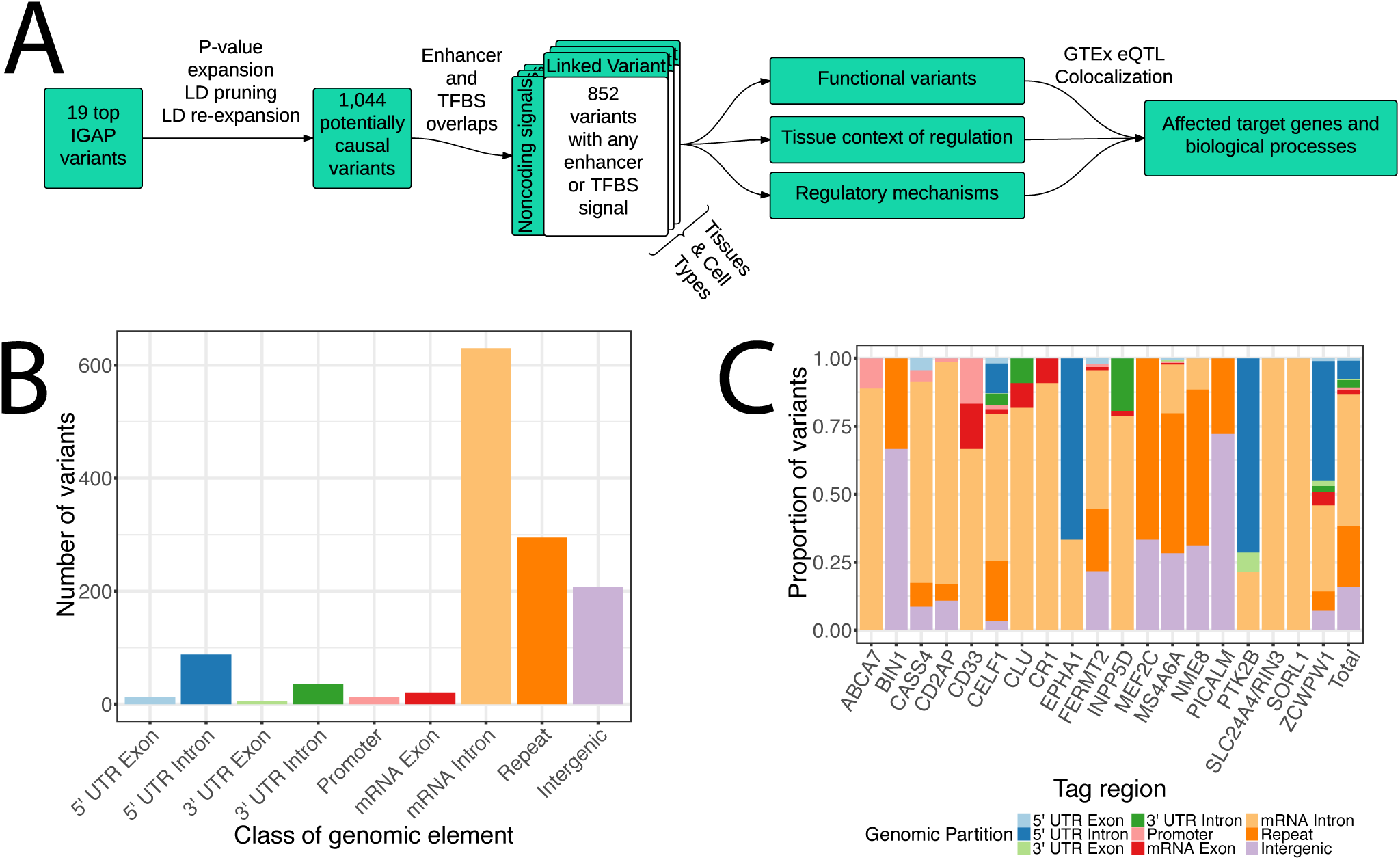
LD expansion and functional annotation of top IGAP hits. a) Flowchart of analysis approach. b) Genomic localization of all variants in *P* value- and LD-expanded set. c) Genomic partition proportions split by tag regions.

## 2. Methods

### 2.1. INFERNO analysis of IGAP top hits

INFERNO (details of the algorithm are described in [23]) was used to analyze 19 top variants from Phase 1 of the IGAP study, excluding the locus near *DSG2* (tagged by rs8093731) which did not replicate in Phase 2 and the *HLA-DRB5* locus (rs9271192) which is difficult to analyze due to the dense LD structure in the major histocompatibility (MHC) region caused by population-specific selective pressure [24]. INFERNO was run using *P* value expansion within one order of magnitude and 500 kilobases (kb) of each tagging variant, and the European population from the 1,000 Genomes Project [25] was used for LD pruning and expansion. For both pruning and expansion, a threshold of r^2^ >= 0.7 was used to define LD blocks. All downstream analyses including lncRNA correlation and pathway analysis were performed as defined in [23].

### 2.2 Luciferase validation

Through molecular cloning techniques, insert sequences including the enhancers (800- 1,739 base pairs (bp) in length, Supplementary Table 1) overlapping each prioritized variant were placed upstream of a minimal promoter and a luciferase reporter gene in a pGL vector. Two different vectors were generated for each prioritized variant: one with the minor allele of the prioritized variant and one with the major allele. Additionally, we generated vectors containing a minimal promoter with no enhancer inserted and another negative control vector with a minimal promoter and a ∼1kb random genomic heterochromatin insert. 300ng of each vector was mixed with one-tenth the amount of a Renilla expressing vector, allowing us to normalize Luciferase expression for transfection efficiency. This mixture was transfected into separate aliquots of K562 cells using the Lonza Nucleofector Device with Kit V. A mock sample was run through the same transfection procedure with no DNA to account for background luminescence. The Promega Dual-Glo system was used to measure Luciferase and Renilla expression. Background-subtracted Luciferase luminescence levels were divided by the corresponding background-subtracted Renilla luminescence, and all ratios were normalized to the average of the minimal promoter condition for quantitative analysis. A total of n = 5 biological replicate experiments were carried out, each including 4 technical replicates per condition.

**Table 1:** IGAP top hits expansion counts and annotation overlaps

| **Tag variant** | **Gene region** | **# *P* value expanded** | **# LD pruned** | **# LD expanded** | **# FANTOM5 overlaps** | **# Roadmap ChromHMM enhancer overlaps** | **# HOMER TFBS overlaps** |
| --- | --- | --- | --- | --- | --- | --- | --- |
| rs4147929 | ABCA7 | 7 | 2 | 9 | 0 | 8 | 6 |
| rs35349669 | BIN1 | 2 | 1 | 3 | 0 | 3 | 1 |
| rs7274581 | CASS4 | 20 | 2 | 23 | 4 | 19 | 10 |
| rs10948363 | CD2AP | 80 | 5 | 83 | 0 | 45 | 30 |
| rs3865444 | CD33 | 3 | 1 | 6 | 0 | 3 | 4 |
| rs10838725 | CELF1 | 92 | 7 | 264 | 13 | 147 | 110 |
| rs28834970 | CLU | 10 | 1 | 11 | 0 | 11 | 7 |
| rs6656401 | CR1 | 20 | 1 | 22 | 0 | 12 | 9 |
| rs11771145 | EPHA1 | 9 | 3 | 9 | 2 | 16 | 2 |
| rs17125944 | FERMT2 | 32 | 7 | 92 | 2 | 79 | 43 |
| rs6733839 | INPP5D | 65 | 2 | 114 | 0 | 66 | 54 |
| rs190982 | MEF2C | 2 | 1 | 3 | 0 | 2 | 1 |
| rs983392 | MS4A6A | 80 | 3 | 173 | 5 | 91 | 75 |
| rs1476679 | TXNDC3 / NME8 | 44 | 5 | 96 | 4 | 44 | 47 |
| rs10792832 | PICALM | 2 | 1 | 18 | 2 | 16 | 11 |
| rs9331896 | PTK2B | 7 | 2 | 14 | 3 | 14 | 8 |
| rs10498633 | SLC24A4 | 5 | 3 | 5 | 4 | 4 | 4 |
| rs11218343 | SORL1 | 1 | 1 | 1 | 0 | 1 | 1 |
| rs2718058 | ZCWPW1 | 15 | 4 | 98 | 0 | 78 | 28 |

Statistical analysis was performed using a linear mixed model treating experimental days as random effects and alleles as fixed effect using the lmerTest package [26] in R v3.4.4 [27]. *P* values for the comparisons between conditions were obtained by analysis of variance (ANOVA) using Satterthwaite’s approximation for degrees of freedom.

## 3. Results

### 3.1 Expansion and annotation of IGAP loci

To identify genetic variants with regulatory potential for LOAD, we used INFERNO to analyze the 19 genome-wide significant loci from Phase I of IGAP (Figure 1a, Table 1) [6]. The region tagged by each top variant is referred to by the name of the nearest gene by convention, although these genes are not necessarily causal for the association signals. For each top variant, we identified all variants within 500kb that had a *P* value within an order of magnitude and the same minor allele effect direction. We pruned these p-value expanded sets by LD into independent variants, which we re-expanded yielding 1,044 unique potentially causal variants (Table 1) for subsequent analyses. These variants were primarily in introns and intergenic regions, with only 17 in mRNA exons (Figure 1b-c).

Next, we overlapped these variants with enhancers defined by bidirectional enhancer RNA (eRNA) transcription in 112 tissues and cell types from the FANTOM5 consortium [17] and by the ChromHMM epigenomic state-based method [28] in 127 tissues and cell types from the Roadmap Epigenomics Project [15,29,30]. This identified 38 variants overlapping FANTOM5 enhancers in 9 tag regions (Table 1). The FANTOM5 tissue with the most enhancer-overlapping variants was monocytes, with 25 overlapping variants, whereas the brain harbored only 6 variants (Supplementary Figure 1a). For the Roadmap data, variants were overlapped with a total of 15 ChromHMM states including 3 types of enhancer states (enhancers, genic enhancers, and bivalent enhancers). 652 unique variants representing all 19 tag regions were found to overlap a ChromHMM-defined enhancer state in at least one tissue (Table 1). Like the FANTOM5 results, primary monocytes from peripheral blood had the most overlapping variants (149 unique variants, Supplementary Figure 1b), but 146 unique variants overlapped enhancers in at least one of the brain-related Roadmap datasets.

We also used INFERNO to find variants affecting transcription factor binding sites (TFBSs) as identified by HOMER [31]. This identified 451 variants representing all 19 tag regions that either increased or decreased TFBS strength (measured by the change in the positional weight matrix, ΔPWM) for 191 unique transcription factors (Supplementary Figure 2). The majority of these overlaps had negative ΔPWM values, reflecting TFBSs disruptions.

### 3.2 Integrative analysis of enhancer enrichment patterns

Using INFERNO’s tissue categorization approach [23] that groups each functional genomics dataset into one of 32 high-level tissue categories, we identified 36 variants from nine tag regions (the *CASS4, CELF1, EPHA1, FERMT2, MS4A6A, NME8, PICALM, PTK2B,* and *SLC24A4*/*RIN3* regions) that overlapped concordant FANTOM5 and Roadmap ChromHMM enhancers in a tissue category (Figure 2a). All of these regions harbored at least one variant with concordant support in the blood category, supporting the hypothesis of immune mechanisms underlying LOAD genetic signals [7–9]. The *CELF1* region was the only one to harbor variants with concordant overlaps of brain enhancers, supporting the unbiased approach of not requiring an *a priori* hypothesis of relevant tissue context. Many variants overlapped FANTOM5 enhancers in the blood category, which included all the immune-related cell lines such as monocytes and macrophages in addition to whole blood (Supplementary Figure 3a). All 22 of the tissue categories sampled by the Roadmap Epigenomics consortium contained ChromHMM-defined enhancer-related states that harbored at least one variant in the expanded set (Supplementary Figure 3b). Again, many variants overlapped Roadmap enhancers in the blood category.

**Figure 2:**
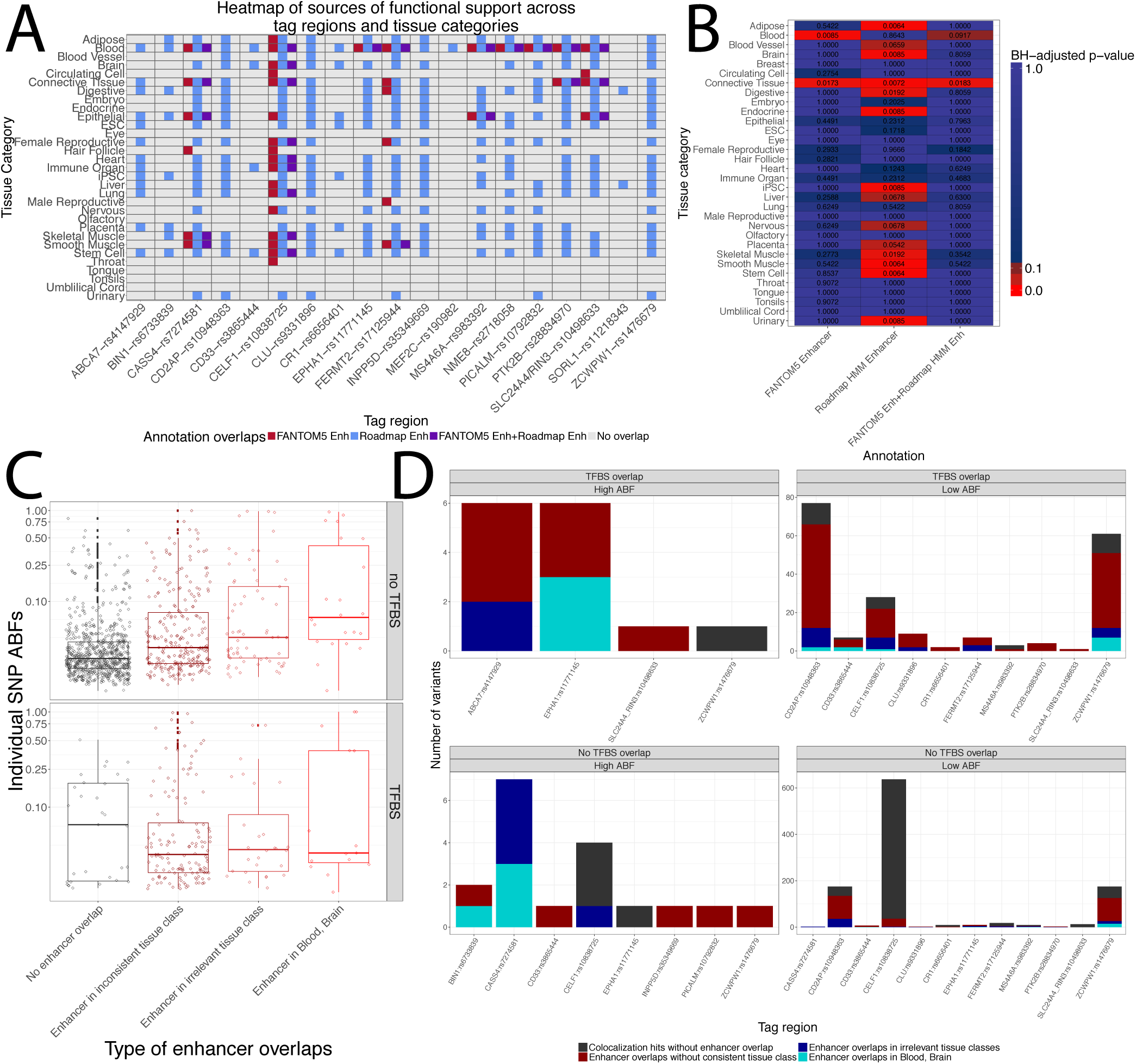
Integrative analysis of annotations for IGAP top hits. a) Integrative tissue context analysis of enhancer overlaps from FANTOM5 and Roadmap datasets. b) Results of LD-collapsed bootstrapping for enhancer annotation overlap enrichments c) Distributions of variant probability of underlying highly colocalized signals stratified by annotation overlap. d) Barplots of numbers of variant – eQTL comparisons across tag regions stratified by motif overlap, enhancer support, and concordant support in a relevant tissue class.

INFERNO includes a method to statistically quantify the enrichments of variants overlapping FANTOM5 enhancers, Roadmap enhancers, or both in each tissue category. This revealed significant enrichments of variants overlapping both FANTOM5 and concordant enhancers in the blood category and a significant enrichment of Roadmap enhancers in the brain category (Figure 2b) as well as several other enrichments including in the connective tissue category, which contains fibroblasts.

### 3.3 Co-localization analysis with GTEx eQTLs

To identify target genes affected by dysregulated enhancers, INFERNO uses expression quantitative trait loci (eQTLs) – variants whose alleles are correlated with differing levels of a target gene – from the Genotype-Tissue Expression (GTEx) project [16] across 44 tissues. Of the 1,044 potentially causal variants, 750 were significant eQTLs in at least one tissue. However, due to dense LD structures in many of our significant regions, this direct overlap approach may yield false positive variants in LD with the truly causal eQTL variant. To address this issue, INFERNO incorporates a Bayesian statistical model (COLOC [32]) to identify co-localized GWAS/eQTL signals with shared causal variants, quantified as the posterior probability P(H_4_). The COLOC method also computes the probability of any individual variant being the shared causal variant, quantified as their Approximate Bayes Factors (ABFs).

We applied COLOC to tissue-specific eQTL signals for 884 unique genes across all 19 tag regions (median number of genes within each region = 34) for 25,435 tests of GWAS – tissue-specific eQTL co-localization (Supplementary Figure 4a, Methods). We identified 153 co-localized GWAS/eQTL signals (P(H_4_) >= 0.5 representing strong support for a shared causal signal [23]) representing 16 tag regions, 37 tissues and 71 target genes (Supplementary Figure 4b). For 32 of these, COLOC identified individual variants with ABF >= 0.5, but in the majority of cases COLOC was not able to prioritize a single causal variant. This is likely caused by dense LD structures where GWAS and eQTL signals are dispersed across all variants in the LD block (Supplementary Figure 4c). Thus, for each co-localized GWAS/eQTL signal we sampled the highest ABF variants until their sum was 0.5 or greater (Supplementary Figure 4d, [23]). Across the 153 co-localized signals, this yielded 1,291 unique variant–tissue–target gene relationships accounted for by 286 unique variants, 182 of which were in the LD-expanded set.

### 3.4 Comparison of enhancer overlaps with eQTL co-localization signals

We next used the INFERNO tissue categorizations to stratify variants in the ABF- expanded sets by whether they affected a TFBS, overlapped any enhancer, and whether the enhancer came from the same tissue category as the eQTL (Figure 2c). For the first stage of variant prioritization, we considered only variants overlapping concordant enhancers, and took two approaches for further prioritization: requiring TFBS overlap (TBFS prioritization) and requiring ABF >= 0.5 (ABF prioritization). TFBS prioritization identified 43 unique variant–tissue–gene sets (20 unique variants across 8 tag regions, Figure 2d, top row) including 15 in the brain or blood categories. ABF prioritization prioritized 14 variant–tissue–gene sets (6 unique variants across 5 tag regions, Figure 2d, left column), including 2 variants which also had motif overlaps. Together, these two approaches identified potentially causal variants in 10 tag regions (Table 2, Supplementary Tables 1-2). We prioritized four of these signals for experimental validation based on prior literature, strength of annotation support, and relevant tissue contexts: *EPHA1, CD33, BIN1*, and *CD2AP*.

**Table 2.** Summary of colocalization results in 4 top prioritized tag regions.

| Tag Region | Top affected mechanism(s) and evidence | Direction of effect | Experimental validation performed? |
| --- | --- | --- | --- |
| *EPHA1* | Whole blood eQTL for ***EPHA1-AS1*** supported by high ABF and increased CEBP TFBS variant rs11765305 | Protective haplotype has strong increase in *EPHA1-AS1* expression | Yes, strong enhancer activity and allelic difference |
| *CD33* | Whole blood eQTL for ***CD33*** with high ABF tag variant rs3865444 | Protective tag variant decreases *CD33* expression | Yes, strong allelic difference |
| *BIN1* | Lymphocyte eQTLs for ***BIN1***, high ABF variant rs4663105 | Risk haplotype decreases *BIN1* expression | Yes, significant allelic difference |
| *CD2AP* | Cerebellar eQTL for ***RP11-385F7.1*** and fibroblast eQTL for ***CD2AP*** with TF disrupting variant rs9367279 | Risk haplotype with moderate, inconsistent effects on lncRNA expression in brain, decrease in *CD2AP* expression | Yes, no K562 enhancer activity |

### *EPHA1* region functional variant upregulates lncRNA affecting the *JAK2* signaling axis

The strongest signal by both annotation and ABF evidence was in the *EPHA1* region, where the variant rs11765305 had an ABF of 0.999 underlying an eQTL for the *EPHA1- AS1* antisense long non-coding RNA (lncRNA) in whole blood (P(H_4_) = 0.516). This variant also colocalized with whole blood eQTLs for the *TAS2R60* taste receptor gene (P(H_4_) = 0.516, ABF = 1.00) and the *TAS2R62P* taste receptor gene (P(H_4_) = 0.537, ABF = 0.714) (Supplementary Table 1). rs11765305 overlapped FANTOM5 and Roadmap enhancers in the blood category, including white blood cells in the myeloid lineage such as monocytes and macrophages (Figure 3a), and creates a stronger binding site for *CEBPB* (ΔPWM score = 1.53), an enhancer-binding transcription factor that is associated with immune-related gene regulation [33]. This increase in TF binding is consistent with the positive effect of the rs11765305 minor allele on *EPHA1-AS1* expression observed in GTEx (β = 1.25, where a ß greater than 1 reflects an increase in gene expression).

**Figure 3:**
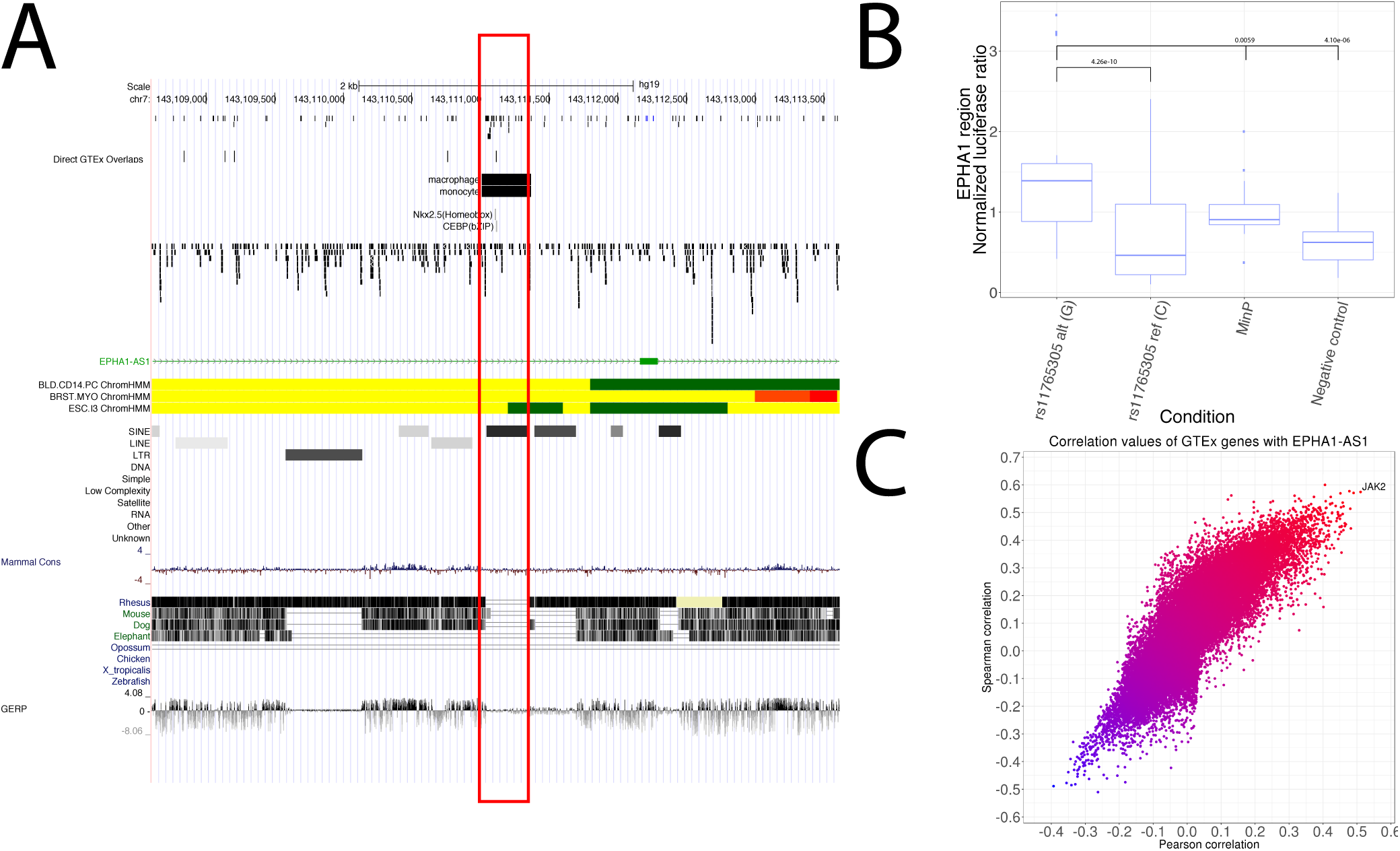
Functional variant in *EPHA1* region upregulates *EPHA1-AS1* lncRNA which regulates the *JAK2* signaling axis. a) Genome browser view of the region around rs11765305 (in red box) including relevant FANTOM5 and Roadmap enhancer annotations. b) Luciferase assay results for rs11765305 in K562 cells. Luciferase expression is normalized against Renilla expression in the same well. Negative control is randomly sampled heterochromatin insert. c) Scatterplot of Pearson and Spearman correlations between expression of *EPHA1-AS1* and all other genes in the genome across all GTEx tissues.

To compare enhancer activity between the major and minor alleles of rs11765305, we performed luciferase assays in K562 leukemia cells, which are from the same myeloid cell lineage as monocytes. Although the major allele had no significant luciferase expression compared to controls, the minor allele had significantly higher expression compared to both controls and the major allele (Figure 3b). These results confirm the predicted monocyte enhancer activity in this region and are consistent with the mechanism that the minor allele of rs11765305 creates a stronger *CEBPB* TFBS, increasing the activity of an enhancer regulating *EPHA1-AS1, TAS2R60,* and *TAS2R62P*.

We next set out to identify the downstream effects of *EPHA1-AS1*, as lncRNAs can modulate gene expression through recruitment of regulatory proteins or binding to target transcripts [34]. INFERNO uses GTEx RNA-seq data to identify genes whose expression is correlated with that of a lncRNA using a threshold of 0.5 on both Pearson and Spearman correlations across 44 tissues [23]. For *EPHA1-AS1*, this yielded one gene, *JAK2* (Pearson r^2^ = 0.517, Spearman r^2^ = 0.582) (Figure 3c). *JAK2* is part of the *JAK2/STAT3* signaling axis, whose disturbance by amyloid-ß leads to memory impairment [35]. The tag variant in this region is protective and rs11765305 has the same effect direction, so INFERNO prioritized a mechanism whereby the protective minor allele of rs11765305 increases *EPHA1-AS1* expression which in turn increases the activity of the *JAK2/STAT3* signaling axis, implying that *JAK2/STAT3* activation may protect against LOAD.

### 3.6 Functional validation of blood regulation of *CD33*

In the *CD33* region, COLOC identified co-localized GWAS/eQTLs for *CD33* itself in whole blood (P(H_4_) = 0.955) and for *AC018755.1* (P(H_4_) = 0.683) in brain hypothalamus. In both cases, rs12459419 was prioritized by concordant enhancer and motif overlap. However, the tag variant rs3865444 had a higher ABF in both cases (0.491 and 0.489, respectively). rs3865444 overlaps Roadmap enhancers in 6 cell lines including primary monocytes and primary T regulatory cells from peripheral blood. In contrast, rs12459419 only overlapped Roadmap enhancers from 3 cell lines including primary T regulatory cells from peripheral blood and fetal brain but lacked the monocyte enhancer overlap (Supplementary Table 1). Additionally, rs3865444 has been extensively studied, with previous work showing that the protective minor allele (A) decreases the levels of *CD33* protein [36], decreases *CD33* mRNA expression consistent with the direction of the GTEx eQTL effect (β = 0.352) [37], and reduces cell surface expression of *CD33* in monocytes [38].

Based on the prior literature, the strong ABF signal, and the monocyte enhancer overlap, we analyzed rs3865444 in our luciferase assays. This found significant increases for the major allele and significant decreases for the minor allele relative to the controls, as well as a striking decrease in enhancer activity of the minor allele relative to the major allele (Figure 4a). This was consistent with prior reports and the GTEx eQTL direction for this variant (ß = 0.352).

**Figure 4:**
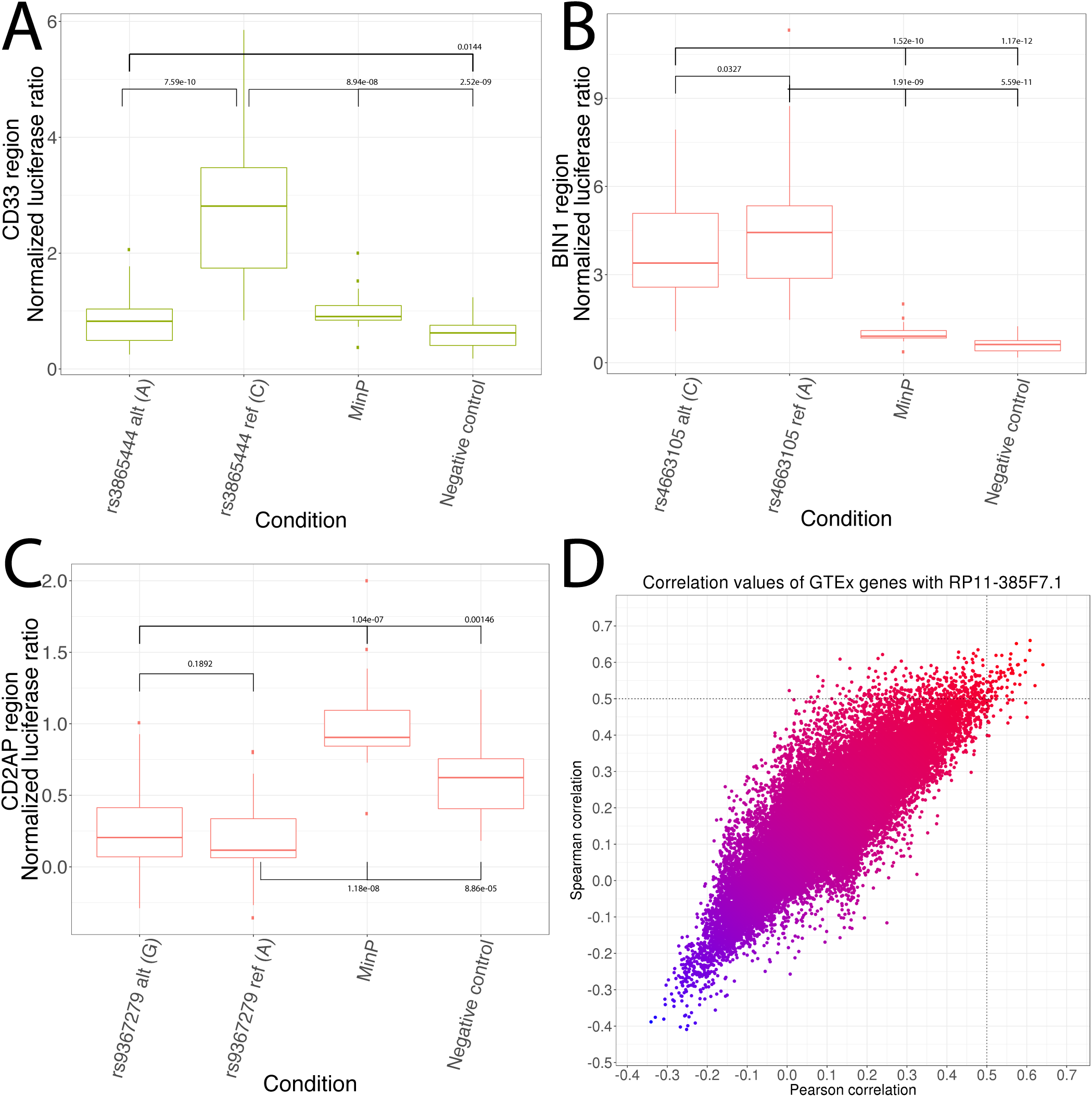
Luciferase and lncRNA analysis in the *BIN1, CD33,* and *CD2AP* regions. a) Luciferase validation in the *CD33* region. b) Luciferase validation in the *BIN1* region. c) Luciferase validation in the *CD2AP* region. d) Scatterplot of Pearson and Spearman correlations between expression of *RP11-385F7.1* (*CD2AP* region) and all other genes in the genome across all GTEx tissues.

### 3.7 Functional validation of lymphocyte regulation of *BIN1*

In the *BIN1* region, INFERNO identified a co-localized GWAS/eQTL for *BIN1* in EBV-transformed lymphocytes (P(H_4_) = 0.652) with the variant rs4663105 prioritized by ABF (ABF = 0.777). This variant overlaps Roadmap enhancers in primary monocyte cells and placenta but does not overlap any TFBSs. rs4663105 has been previously associated with LOAD risk, and an insertion in that region was associated with increased *BIN1* expression [39]. This previous study found no difference in luciferase activity between the two alleles of rs4663105 in SKNSH-SY5Y and HEK cells. However, their construct only spanned 60bp around the variant, whereas the monocyte enhancer is 800bp. Therefore, we cloned the full Roadmap enhancer region (Supplementary Table 1) for luciferase assays in K562 cells, which are more relevant to the functional annotations in this region. This found significantly increased enhancer activity for both alleles of rs4663105 relative to the control vectors, and a slight but significant decrease of the minor allele relative to the major allele (p = 0.0328, Figure 4b), consistent with the direction of the GTEx eQTL (ß = 0.496).

### 3.8. *CD2AP* region variants modulate lncRNA with widespread brain regulatory effects

Finally, in the *CD2AP* region, INFERNO prioritized several co-localized signals including *RP11-385F7.1* in brain cerebellar hemisphere and cerebellum (P(H_4_) = 0.904 and 0.923, respectively) and an eQTL for *CD2AP* in fibroblasts (P(H_4_) = 0.801). TFBS prioritization implicated rs9367279, which overlaps Roadmap enhancers in 33 tissues/cell lines from 13 tissue categories and disrupts a CArG-box binding site (ΔPWM = -1.38) for the MADS-box family of transcription factors, which includes the enhancer-related factors *SRF* and *MEF2A* [40,41]. *CD2AP* encodes a scaffolding molecule that regulates the actin cytoskeleton and is involved in endocytic processes [42], while *RP11-385F7.1* is a lncRNA near the promoter for the *CD2AP* gene.

We performed luciferase assays, but both alleles of rs9367279 had significantly decreased enhancer activity relative to the controls, and there was no strong difference between the two alleles, suggesting that this enhancer may not be active in K562 cells (Figure 4c, p = 0.1892). *RP11-385F7.1* was strongly correlated with 64 transcripts (Figure 4d, Supplementary Table 4), and we performed pathway analysis of these targets using the WebGestalt tool [43,44] to interpret the increased number of targets relative to section 3.5, but found no enrichments after controlling for false discovery rate. The gene with the strongest Pearson correlation was *PPP1R16A* (Pearson r^2^ = 0.641, Spearman r^2^ = 0.593) and the gene with the strongest Spearman correlation was *COQ4* (Pearson r^2^ = 0.608, Spearman r^2^ = 0.660). *PPP1R16A*, also known as *MYPT3*, directs the protein phosphatase PP1c to its targets and is involved in actin binding and G-protein coupled receptor pathways [45]. *COQ4* is part of the coenzyme Q biosynthesis pathway, an antioxidant that may modify LOAD-associated oxidative damage [46]. The eQTL effects of rs9367279 on *RP11-385F7.1* are weak and inconsistent between the two brain regions (ß = 0.969 in cerebellar hemisphere and 1.194 in cerebellum), suggesting that rs9367279 contributes to fine-scale regulation of *RP11-385F7.1* in brain, although it has a relatively strong repressive effect on *CD2AP* in fibroblasts (ß = 0.505).

## 4. Discussion

Our application of INFERNO to LOAD GWAS data prioritized perturbations of tissue-specific regulatory mechanisms in 10 IGAP tag regions (Table 2, Supplementary Table 3). In the *EPHA1, CD2AP, CELF1*, and *CASS4* regions, the target genes of the co-localized GWAS/eQTL signals included lncRNAs, so identifying affected enhancers and target genes may be only the first step towards understanding genetic effects on regulatory networks contributing to disease pathogenesis. The tissue classification approach implemented in INFERNO also enabled the unbiased investigation of the relevant tissue contexts affected by each genetic signal. Limiting our analysis to only brain datasets would have missed the blood-category signals that we detected. These immunity-related signals are in line with other recent work highlighting neuroinflammation as a crucial component of LOAD pathogenesis and etiology [7–9]. INFERNO did not identify regulatory mechanisms in all 19 of the IGAP regions, and this may be driven by several aspects of this analysis. We are limited by the sample sizes and sets of tissues that were assayed by the FANTOM5, Roadmap, and GTEx consortia and the number of datasets that went into each tissue category, with some categories being much more sparsely sampled than others [23]. Another consideration is that this regulatory analysis focused on transcriptional enhancers, but it is possible that the noncoding signals in the unexplained tag regions affect other regulatory mechanisms such as small noncoding RNA (sncRNA) loci. Previous studies implicated sncRNA dysregulation in LOAD pathogenesis [47], suggesting that this will be a fruitful approach for future analysis efforts.

In conclusion, our application of INFERNO to IGAP GWAS data yielded insights into the regulatory mechanisms affected by noncoding LOAD-associated genetic variants. Experimental validation supported our computationally predicted regulatory effects, suggesting that our approach is able to prioritize truly causal regulatory mechanisms at GWAS loci for post-GWAS experiments. Incorporating more functional genomics data as it is generated in concert with more refined validation experiments using a broader range of cell types and molecular techniques will yield insights into a range of phenotypes.

## Figure legends and tables

**Supplementary Figure 1: Number of enhancer overlaps across individual tissues from FANTOM5 and Roadmap.** a) Number of variants overlapping eRNA-defined enhancers across 112 FANTOM5 tissue and cell type facets. b) Number of variants overlapping each type of ChromHMM-defined enhancer state across 127 Roadmap tissues and cell types.

**Supplementary Figure 2: HOMER motif overlap ΔPWM distributions**

**Supplementary Figure 3: Number of enhancer overlaps across tissue categories sampled in FANTOM5 and Roadmap.** a) Number of variants overlapping eRNA-defined enhancers in each tag region across the 28 tissue categories that include FANTOM5 samples. b) Number of variants overlapping any of the three ChromHMM-defined enhancer states in each tag region across the 22 tissue categories that include Roadmap samples.

**Supplementary Figure 4:Colocalization analysis of GTEx eQTLs with IGAP GWAS signals.** a) Distributions of the 5 colocalization hypotheses across all tissues and tag regions. b) Histograms of P(H_4_) for highly colocalized (P(H_4_) >= 0.5) signals across tag regions. c) Histograms of the approximate Bayes factor values of the most supported variants across tag regions. d) Histograms of the number of variants required to cumulatively account for 50% of the individual variant probability (ABFs) for each colocalization signal across tag regions.

**Supplementary Table 1: Luciferase enhancer regions** (Excel file)

**Supplementary Table 2: Full colocalization annotation results. Top signals in each region are highlighted.** (Excel file)

**Supplementary Table 3: Summary of colocalization results in 6 non-prioritized regions**

**Supplementary Table 4.** Highly correlated genes with *RP11-385F7.1*. (Excel table)

| Tag Region | Top affected mechanism(s) and evidence | Direction of effect | Experimental validation performed? |
| --- | --- | --- | --- |
| *ABCA7* | Digestive system eQTLs for ***ABCA7***, high ABF variant rs4147929 | Risk haplotype increases *ABCA7* expression | No, irrelevant tissue category |
| *CASS4* | Fibroblast eQTL for ***CASS4*** with high ABF variant rs6014724, blood eQTL for ***CASS4*** with high ABF variant rs927174 | Protective haplotype increases expression in fibroblasts, lowers in blood | No, inconsistent effect directions and lack of TFBS overlap |
| *CELF1* | Brain eQTL for ***RP11-750H9.5*** supported by rs7947450 with moderate TF disruption, fibroblast eQTL for ***MADD*** with high ABF variant rs11039281 | Risk haplotype decreases expression of eQTL genes | No, very dense LD region, molecular cloning for single variant failed |
| *CLU* | Epithelial and digestive eQTLs for **ZNF395** and **FZD3**, both supported by rs2070926 with TFBS overlap | Protective haplotype decreases *ZNF395* and *FZD3* expression | No, irrelevant tissue categories |
| *FERMT2* | Skeletal muscle eQTL for **FERMT2**, several variants with enhancer + motif support | Risk haplotype decreases *FERMT2* expression | No, irrelevant tissue |
| *ZCWPW1* | Brain eQTLs in several different regions for **PVRIG** and **STAG3** supported by rs1727138 with strong TFBS disruption | rs1727138 has inconsistent effects on expression levels across tissues & genes | No, inconsistent effect directions for rs1727138 and its GWAS effect direction does not match the tag variant |

## Author contributions

- Computational analysis: AAW, MT
- Validation experiments: AAW, MT, JW, BD, NV, MJ, YFC, YZ, AK
- Analysis design: AAW, MT, CDB, LSW, GDS
- Writing and editing: AAW, MT, NV, EEM, YYL, CDB, LSW, GDS

## Acknowledgements

- This work was supported by the National Institutes of Health [grant numbers U01-AG032984, UF1-AG047133, U54-AG052427, U24-AG041689, R01- GM099962, P30-AG010124, RF1-AG055477, U54-NS100693, and T32-AG00255].
- We thank the International Genomics of Alzheimer’s Project (IGAP) for providing summary results data for these analyses. The investigators within IGAP contributed to the design and implementation of IGAP and/or provided data but did not participate in analysis or writing of this report. IGAP was made possible by the generous participation of the control subjects, the patients, and their families. The i–Select chips was funded by the French National Foundation on Alzheimer’s disease and related disorders. EADI was supported by the LABEX (laboratory of excellence program investment for the future) DISTALZ grant, Inserm, Institut Pasteur de Lille, Université de Lille 2 and the Lille University Hospital. GERAD was supported by the Medical Research Council (Grant n° 503480), Alzheimer’s Research UK (Grant n° 503176), the Wellcome Trust (Grant n° 082604/2/07/Z) and German Federal Ministry of Education and Research (BMBF): Competence Network Dementia (CND) grant n° 01GI0102, 01GI0711, 01GI0420. CHARGE was partly supported by the NIH/NIA grant R01 AG033193 and the NIA AG081220 and AGES contract N01–AG–12100, the NHLBI grant R01 HL105756, the Icelandic Heart Association, and the Erasmus Medical Center and Erasmus University. ADGC was supported by the NIH/NIA grants: U01 AG032984, U24 AG021886, U01 AG016976, and the Alzheimer’s Association grant ADGC–10–196728.
- We gratefully acknowledge Adam Naj, Ian Mellis, and Eddie Lee for providing writing help and feedback on these results.

